# SEX-DEPENDENT MODULATION OF BEHAVIORAL ALLOCATION VIA VENTRAL TEGMENTAL AREA-NUCLEUS ACCUMBENS SHELL CIRCUITRY

**DOI:** 10.1101/2025.02.09.637318

**Authors:** Kristen A. McLaurin, Jessica M. Illenberger, Hailong Li, Rosemarie M. Booze, Charles F. Mactutus

## Abstract

Diagnostic criteria for substance use disorder, cocaine type (i.e., cocaine use disorder), outlined in the 5^th^ edition of the *Diagnostic and Statistical Manual,* imply that the disorder arises, at least in part, from the maladaptive allocation of behavior to drug use. To date, however, the neural circuits involved in the allocation of behavior have not been systematically evaluated. Herein, a chemogenetics approach (i.e., designer receptors exclusively activated by designer drugs (DREADDs)) was utilized in combination with a concurrent choice self-administration experimental paradigm to evaluate the role of the mesolimbic neurocircuit in the allocation of behavior. Pharmacological activation of hM3D(G_q_) DREADDs in neurons projecting from the ventral tegmental area (VTA) to the nucleus accumbens (AcbSh) induced a sex-dependent shift in the allocation of behavior in rodents transduced with DREADDs. Specifically, male DREADDs animals exhibited a robust increase in responding for a natural (i.e., sucrose) reward following pharmacological activation of the VTA-AcbSh circuit; female DREADDs rodents, in sharp contrast, displayed a prominent decrease in drug-reinforced (i.e., cocaine) responding. The sequential activation of hM3D(G_q_) and KORD DREADDs within the same neuronal population validated the role of the VTA-AcbSh circuit in reinforced responding for concurrently available natural and drug rewards. Collectively, the VTA-AcbSh circuit is fundamentally involved in behavioral allocation affording a key target for the development of novel pharmacotherapies.

## INTRODUCTION

In 2023, over 1.25 million individuals living in the United States met the *Diagnostic and Statistical Manual of Mental Disorders* (5^th^ ed.; DSM-5) criteria for substance use disorder, cocaine type (i.e., cocaine use disorder; [1]). The DSM-5 outlines 11 diagnostic criteria for cocaine use disorder, whereby an individual meeting at least two criteria within a 12-month period exhibits a problematic pattern of use [2]. Fundamentally, six of the 11 DSM-5 diagnostic criteria tap the allocation of behavior towards the pursuit and use of drug reinforcers over alternative reinforcers. The DSM-5, therefore, utilizes a behavioral-centric perspective for the diagnosis of cocaine use disorder, thereby implying cocaine use disorder arises, at least in part, from the maladaptive allocation of behavior to drug use (for review, [3]). Nonpharmacological treatment strategies afford additional support for the behavioral-centric perspective, whereby behavioral interventions that reallocate behavior to alternative reinforcers have successfully decreased substance use [4–6]. To date, however, the neural circuits involved in the allocation of behavior have not been fully elucidated; establishing the neural circuits that specify behavioral allocation may provide a key target for the development of novel pharmacotherapies.

Chemogenetics encompasses the process of genetically engineering proteins (e.g., G Protein-Coupled Receptors (GPCRs): [7–8]; Ligand-Gated Ion Channels: [9–10]; Kinases: [11–12]) to respond to otherwise inert small molecules. The superfamily of GPCRs, in particular, represents an attractive engineering target, as they constitute the largest class of cell surface receptors, recognize a variety of ligands, and are expressed in most neuronal cells. Structurally, GPCRs are characterized by seven hydrophobic transmembrane α helices and three intracellular and extracellular loops with an amino and carboxyl terminus, respectively [13–14]. From a mechanistic perspective, activation of a GPCR by an extracellular ligand triggers a prototypical conformational change in the receptor facilitating the interaction with a heterotrimeric G protein that consists of three distinct subunits (i.e., α, β, and γ). The α subunits of G proteins confer, at least in part, the specificity of functional activity, whereby the G_s_ and G_i_ alpha subunits are involved in the stimulation [15] and inhibition [16] of adenylate cyclase, respectively; the G_q_ alpha subunit activates phospholipase C [17]. Thus, the development of a family of GPCRs that are selectively activated by a pharmacologically inert compound has the potential to revolutionize our approach to dissecting the neural circuits involved in behavior.

Indeed, in 2007, designer receptors exclusively activated by designer drugs (DREADDs), the third generation of GPCR-based chemogenetic tools, were developed. Specifically, a directed molecular evolution approach was utilized to engineer a family of human muscarinic GPCRs (i.e., hM1D_q_, hM2D_i_, hM3D_q_, hM4D_i_, hM5D_q_) that are selectively activated by highly specific chemical actuators (e.g., Clozapine-*N*-Oxide (CNO): [8]; Compound 21 (C21): [18–19]). In alignment with the specificity of functional activity conferred by the α subunits of G proteins, G_q_ and G_i_-DREADDs activate [20] and inhibit [8] neuronal activity, respectively.

Nevertheless, both G_q_ and G_i_-DREADDs respond to the same pharmacologically inert ligand (i.e., CNO or C21), thereby precluding the bidirectional control of neuronal activity. Given these limitations, Vardy et al. [21] evolved the human Κ-opioid receptor (KOR) to respond to Salvinorin B (SalB; [21]), another highly specific chemical actuator [21–22] that exhibits rapid uptake and clearance in the brain [23]. Co-expression of G_q_-based DREADDs and KOR-based DREADDs, therefore, affords a unique opportunity to control neuronal activity in a sequential and bidirectional manner for the interrogation of neural circuits underlying the allocation of behavior.

Behavioral allocation, which requires an organism to assign value to the available options, may be mediated, at least in part, by mesolimbic neurocircuitry. The mesolimbic pathway is comprised of the ventral tegmental area (VTA) and nucleus accumbens (AcbSh), whereby the AcbSh is innervated by excitatory dopaminergic and glutamatergic projections, as well as inhibitory gamma-aminobutyric acid afferents, from the VTA. Indeed, the striatum is fundamentally involved in evaluating the relative value [24], magnitude (e.g., one versus three food pellets; [25–26]), and probability [25] of the available outcomes. More specifically, accumbal dopamine (DA) appears highly involved in behavioral allocation, as DA depletion in the AcbSh alters choice selection dependent upon the effort requirements of the response [27–28]. Measurement of dopaminergic neurons in the mesolimbic pathway, however, affords direct evidence for the role of mesolimbic neurocircuitry in behavioral allocation, whereby the activity of DA neurons encodes both extrinsic (e.g., Probability: [29]; Delay: [30]) and intrinsic [31] factors of reward value. Nevertheless, prior studies are limited by the inclusion of only one reinforcer type (i.e., Non-Drug Reinforcers).

Thus, the present study was undertaken to establish the role of mesolimbic neurocircuitry in drug-biased choice. In preclinical biological systems, a concurrent choice self-administration experimental paradigm affords an opportunity to delineate the relative reinforcing effects of a drug (e.g., cocaine) in comparison to an alternative reinforcer (e.g., sucrose; for review, [32]). After establishing a history of sucrose and cocaine self-administration, rodents were trained and assessed in a concurrent choice self-administration experimental paradigm; the subsequent transduction with DREADDs or vehicle and treatment with C21 and/or SalB afforded an opportunity to evaluate the role of mesolimbic neurocircuitry in the allocation of behavior. Establishing the neural circuits that specify behavioral allocation may provide a key target for the development of novel pharmacotherapies.

## MATERIALS AND METHODS

### Experimental Design

A within-subjects repeated-measures experimental design was utilized to establish the role of dopaminergic VTA neurons projecting to the AcbSh in natural (sucrose) and drug (cocaine) reinforcement (**Figure 1**).

**Figure 1.**
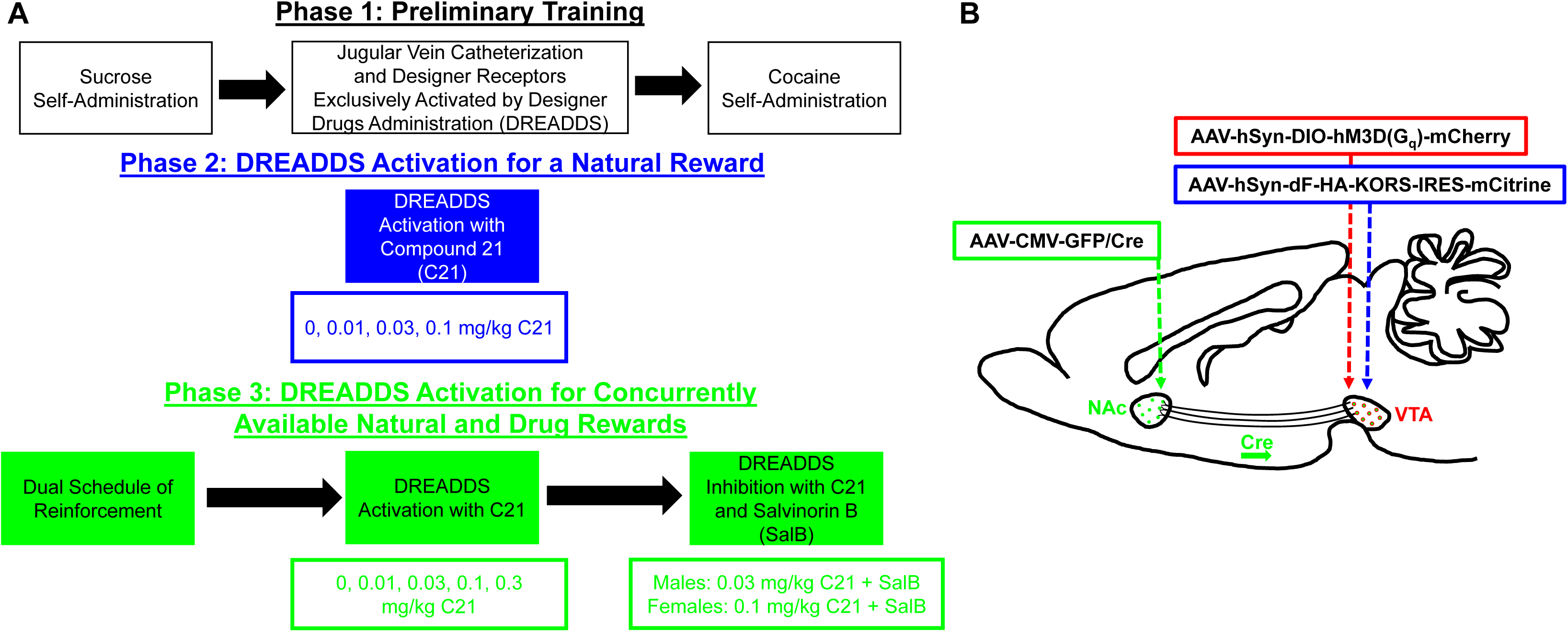
Experimental Design. (**A**) A schematic of the experimental design. (**B**) Three viral vectors were administered to DREADDs transduced rodents. AAV-CMV-GFP/Cre was injected into the nucleus accumbens (AcbSh) of male and female Fischer F344/N rats, whereas AAV-hSyn-DIO-hM3D(G_q_)-mCherry and AAV-hSyn-dF-HA-KORS-IRES-mCitrine were administered into the ventral tegmental area (VTA). AAV-CMV-GFP/Cre is retrogradely transported along the axons of neurons in the VTA to the soma where it reorients either AAV-hSyn-DIO-hM3D(G_q_)- mCherry or AAV-hSyn-dF-HA-KORS-IRES-mCitrine. Sham animals were transduced with AAV-CMV-GFP/Cre in the AcbSh and vehicle in the VTA.

Adult intact male (*n*=20) and female (*n*=20) Fischer F344/N rats were acquired from Envigo Laboratories (Indianapolis, IN). To preclude violation of the independence assumption inherent in many traditional statistical approaches (e.g., analysis of variance (ANOVA); [33–34]), unrelated animals (i.e., no more than one male and one female from each litter) were requested from the supplier. After the completion of Sucrose Self-Administration, rodents were randomly assigned to receive either bilateral DREADDs (i.e., VTA: pAAV-hSyn-DIO-hM3D(G_q_)-mCherry, pAAV-hSyn-dF-HA-KORD-IRES-mCitrine; AcbSh: AAV-CMV-GFP/Cre) or sham (i.e., VTA: Vehicle; AcbSh: AAV-CMV-GFP/Cre) infusions, yielding sample sizes of DREADDs, *n*=24 (Male: *n*=12, Female: *n*=12) and Sham, *n*=16 (Male: *n*=8, Female: *n*=8). Appropriate sample sizes for statistical power of 0.80 were established *a priori* using statistical power analyses (G*Power 3, Version 3.9.9.7; [35]), whereby estimates were based on an effect size of 0.30 and an alpha (α) value of 0.05.

Between-subject factors included biological sex (Male vs. Female) and DREADDs infusion (DREADDs vs. Sham). Within-subject factors included dose (C21: 0, 0.01, 0.03, 0.1, 0.3 mg/kg; SalB: 0.15 mg/kg), time, and reinforcer (Sucrose vs. Cocaine), as appropriate. Given the implementation of highly complex and heterogeneous statistical analyses, additional details are reported in the Statistical Analysis section below.

### Animals

Upon delivery to the animal vivarium, rodents were pair- or group-housed with animals of the same sex. After jugular vein catheter implantation and stereotaxic infusion of viral vectors, rodents were single-housed for the remainder of experimentation. Rats were provided with *ad libitum* access to rodent food (Pro-Lab Rat, Mouse, Hamster Chow #3000) and water throughout the study unless otherwise specified. The animal vivarium was maintained at approximately 20 ± 2°C, 50 ± 10% relative humidity with a 12-hour light/12-hour dark cycle (lights on at 7:00h and lights off at 19:00h).

Rodents were housed in AAALAC-accredited facilities at the University of South Carolina (USC; Federal Assurance #D16-00028) using the guidelines established by the National Institutes of Health in the Guide for the Care and Use of Laboratory Animals. The USC Institutional Animal Care and Use Committee approved the project protocol.

### Apparatus

Assessments were conducted in twenty operant chambers (ENV-008; Med-Associates, St. Albans, VT) enclosed within a sound-attenuating cabinet and controlled by Med-PC computer interface software. A 5 cm x 5 cm opening (ENV 202M-S) on the front panel of the operant chamber afforded access to a recessed 0.01 cc dipper cup (ENV-202C) containing the sucrose solution. Two retractable “active” metal levers (ENV-112BM) were also located on the front panel of the operant chamber. The rear panel of the operant chamber contained one non-retractable “inactive” metal lever and a 28-V house light; the house light turned off following responses on the “active” lever during Cocaine Self-Administration and Concurrent Choice Self-Administration. Reinforcement was delivered after the rodent responded on the “active,” but not “inactive,” lever.

Intravenous (IV) cocaine infusions were delivered through a water-tight swivel (Instech 375/22ps 22GA; Instech Laboratories, Inc., Plymouth Meeting, PA) using a syringe pump (PHM- 100). Tygon tubing (ID, 0.020 IN; OD, 0.060 IN) enclosed by a stainless-steel tether (Camcaths, Cambridgeshire, UK) was utilized to connect the water-tight swivel to the back mount of the animal. A Med-PC computer program used an animal’s daily body weight to calculate pump infusion times.

### Phase 1: Preliminary Training

#### Sucrose Self-Administration

*Autoshaping.* Following the completion of magazine training, which was conducted using the methods outlined in Lacy et al. [36], rodents were subsequently trained to lever-press for a 5% *w/v* sucrose solution on a fixed-ratio 1 (FR(1)) schedule of reinforcement. During the 42-min training session, rodents were reinforced after responding on either of the “active” levers; a non-contingent reinforcer was also delivered every 10 min. Nevertheless, with regards to contingent reinforcement, an animal was limited to five consecutive presses on a single “active” lever and a maximum of 120 reinforcers. Successful acquisition of the autoshaping procedure required animals to achieve at least 60 reinforcers for three consecutive days.

The majority of animals (Female: *n*=20; Male: *n*=16) successfully acquired the autoshaping task within 60 training sessions. Animals that failed to meet the criteria after 60 training sessions were water-restricted for up to 18 hours; *ad libitium* access to water was provided after rodents met the criteria under water-restricted conditions. To ensure autoshaping proficiency, animals were required to continue training under non-restricted conditions until meeting criteria. A small subset (Male: *n*=3) of rodents failed to successfully acquire the autoshaping procedure within 110 training sessions, necessitating an increased sucrose solution concentration (i.e., 10% *w/v* instead of 5% *w/v*).

To ensure sufficient sucrose self-administration training, rodents also completed FR(1) training, as well as sucrose dose-response assessments under progressive and fixed-ratio schedules of reinforcement. Detailed methodology for dose-response evaluations has been previously reported [37].

#### Intravenous Catheterizations and Stereotaxic Surgery

After the completion of Sucrose Self-Administration, rodents underwent surgery for two complementary purposes, including 1) to bilaterally infuse DREADDs into the brain; and 2) to implant an IV catheter. Inhalant sevoflurane was utilized to induce (5-7%) and maintain (3-4%) anesthesia throughout the surgical procedure.

First, using stereotaxic procedures, rodents were bilaterally infused with either DREADDs (i.e., VTA: pAAV-hSyn-DIO-hM3D(G_q_)-mCherry, pAAV-hSyn-dF-HA-KORD-IRES-mCitrine; AcbSh: AAV-CMV-GFP/Cre) or sham (i.e., VTA: Vehicle; AcbSh: AAV-CMV-GFP/Cre). To selectively stimulate neurons of the VTA that project to the AcbSh, a “retro-DREADD” [38] technique was implemented whereby Cre-recombinase and FLEX-DREADD constructs were utilized to restrict expression of DREADDs in VTA neurons that project to the AcbSh.

Fundamentally, a standard Cre-recombinase-dependent adeno-associated virus was utilized to create both the hM3D(G_q_) and KORD DREADDs utilized in the present experiment.

Animals were placed in a stereotaxic apparatus (Model 900; Kopf Instruments, Tujunga, CA) and the scalp was exposed. For infusions of AAV-CMV-GFP/Cre into the AcbSh, small holes (0.4 mm) were drilled bilaterally 7 mm into the skull at 0.5 mm and 1.2 mm lateral and rostral to Bregma, respectively. For infusions of DREADD vectors or vehicle into the VTA, 0.4 mm holes were drilled bilaterally 8 mm into the skull at 1 mm and 5 mm lateral and caudal to Bregma, respectively.

Second, using the methods reported by Bertrand et al. [39], rodents were implanted with a chronic indwelling jugular catheter. Specifically, a sterile IV catheter was implanted into the right jugular vein and secured with sterile sutures (4-0 Perma-Hand Silk; EthiconEnd-Surgery, Inc., Cincinnati, OH). The back mount of the catheter was connected to an acrylic pedestal embedded with mesh and subcutaneously implanted on the dorsal surface of the scapulae; sutures (4-0 Monoweb) were utilized to stitch the back mount into place.

Butorphanol (Dorolex; 1.0 mg/kg; Merck Animal Health, Millsboro, DE) and the antibiotic gentamicin sulfate (1%; 0.2 mL; VEDCO, Saint Joseph, MO) were administered subcutaneously and intravenously, respectively, immediately after surgery. Rodents were monitored in a heat-regulated warm chamber and returned to the colony room after recovery from anesthesia.

For 10 days after surgery, catheters were flushed with a post-flush solution containing heparin, an anti-coagulant (2.5%; APP Pharmaceuticals, Schaumburg, IL), and gentamicin sulfate (1%), an antibiotic. Rodents began cocaine self-administration four days after jugular vein catheterization. Jugular vein catheters were flushed prior to and after operant testing each day with a 0.9% saline solution (Baxter, Deerfield, IL) and post-flush solution (i.e., 2.5% heparin and 1% gentamicin sulfate), respectively.

#### Cocaine Self-Administration

*Fixed-Ratio 1 Schedule of Reinforcement.* Rodents responded on one of two “active” levers for 0.2 mg/kg/infusion of IV cocaine. Once animals successfully earned their first cocaine infusion, 60-min assessments were conducted for five days on an FR(1) schedule of reinforcement.

*Progressive Ratio Schedule of Reinforcement.* Subsequently, rodents responded for 0.75 mg/kg/infusion of cocaine according to a PR schedule of reinforcement, whereby the response requirement was increased progressively according to the following exponential function [5e^(reinforcer^ ^number^ ^x^ ^0.2)^]-5 [40]. For seven days, 120-min PR assessments were conducted.

### Phase 2: Designer Receptors Exclusively Activated by Designer Drugs Activation for a Natural Reward

Rodents were intravenously injected with C21 (0, 0.01, 0.03, or 0.1 mg/kg; adjusted from [41]), a DREADDs agonist, to evaluate the role of neurons in the VTA-AcbSh circuit in natural (i.e., sucrose) reward responding on an FR(1) schedule of reinforcement. First, rodents were treated intravenously with 1.0 mL/kg saline for two days to establish baseline sucrose-maintained responding. Subsequently, on testing days, which occurred every other day, rodents were intravenously injected with doses of C21 in an ascending fashion. Maintenance days, whereby animals were treated intravenously with 1.0 mL/kg saline, occurred on the intervening non-test days.

### Phase 3: Designer Receptors Exclusively Activated by Designer Drugs Activation for Concurrently Available Natural and Drug Rewards

#### Concurrent Choice Self-Administration

After establishing both sucrose and cocaine self-administration, rodents were assessed in a concurrent choice self-administration experimental paradigm for seven days. During a 60- min test session, animals responded on an FR(1) schedule of reinforcement for either 5% (*w/v*) sucrose or 0.2 mg/kg/infusion of IV cocaine. The position (i.e., left or right) of the sucrose-paired lever was balanced between groups.

The test session began with four forced-choice trials for two sucrose and two cocaine reinforcers; only one “active” lever was presented to the animal during the forced-choice trials. Subsequently, the presentation of both “active” levers allowed the rodents to freely choose between sucrose and cocaine. A 20-sec time-out (i.e., retraction of the “active” levers and the house light turned off) occurred after every response.

#### Designer Receptors Exclusively Activated by Designer Drugs Activation with Compound 21

Subsequently, male and female rats were intravenously injected with C21 (0, 0.01, 0.03, 0.1, or 0.3 mg/kg), a DREADDs agonist, to evaluate the role of neurons in the VTA-AcbSh circuit in concurrent choice self-administration on an FR(1) schedule of reinforcement. After one maintenance session (i.e., IV treatment with 1.0 mL/kg saline), animals were intravenously injected with C21 prior to beginning the 60-min test session. Doses of IV C21 were administered on test days in ascending order. On intervening non-test days, rodents completed a maintenance session.

### Phase 4: Designer Receptors Exclusively Activated by Designer Drugs Inhibition for Concurrently Available Natural and Drug Rewards

Finally, to determine if the effects of C21 on concurrently available natural and drug rewards are bidirectional, IV C21 inoculation was followed by IV Sal B (0.15 mg/kg; adjusted from [21]). Administration of C21 prior to the concurrent choice self-administration experimental paradigm significantly altered responding in a sex-dependent manner (See Results Below).

Therefore, the 0.03 mg/kg dose of C21 was administered prior to Sal B in male animals, whereas female rodents were treated with 0.1 mg/kg C21 prior to Sal B.

Rodents were treated with C21 and returned to their home cages for approximately 15 min, at which time they were intravenously injected with Sal B. After inoculation with Sal B, the concurrent choice self-administration experimental paradigm was initiated.

### Phase 5: Verification of Cannula Placement and Designer Receptors Exclusively Activated by Designer Drugs Expression

Following the completion of behavioral assessments, cannula placement (DREADDs: *n*=16; Sham: *n*=16) and DREADDs expression (DREADDs: *n*=7; Sham: *n*=2) was confirmed in at least a subset of animals. Rodents were deeply anesthetized with 5% sevoflurane (Abbot Laboratories, North Chicago, IL) and humanely sacrificed via transcardial perfusion (100 mL of 100 mM phosphate buffered saline followed by 150 mL of 4% paraformaldehyde). After removal, the rat brain was post-fixed in 4% paraformaldehyde before being sliced coronally (100 µM).

Coronal brain slides were mounted on microscope slides (Superfrost Plus; Fisher Scientific, Hampton, NH) using Cytoseal XYL (Thermo Fisher Scientific, Waltham, MA). Z-stack images were obtained using a confocal microscopy system (Nikon TE-2000E and Nikon’s EZ-C1 Software (version 3.81b).

### Statistical Analysis

Analysis of variance (ANOVA) and non-linear regression approaches were utilized to statistically analyze all data (GraphPad Prism 10 Software, Inc., La Jolla, CA; SAS/STAT Software 9.4, SAS Institute, Inc., Cary, NC). A *p* value of 0.05 was established for statistical significance.

Preliminary training, including Sucrose Self-Administration and Cocaine Self-Administration, was analyzed using non-linear regression (GraphPad Prism 10 Software, Inc.) and/or generalized linear mixed effects models (GLMM; SAS/STAT Software 9.4). With regards to Sucrose Self-Administration, the number of days to criteria served as the dependent variable of interest. Some animals died after the completion of Sucrose Self-Administration, but before the initiation and/or completion of Cocaine Self Administration (DREADD Male, *n*=1; DREADD Female, *n*=2; Sham Male, *n*=1; Sham Female, *n*=1); data collected from these animals was included through their final assessment. For Cocaine Self-Administration, a GLMM was utilized to analyze the number of cocaine reinforcers earned while responding under a progressive ratio schedule of reinforcement, whereby day served as a within-subjects factor. The factor of biological sex served as a between-subjects factor during preliminary training.

The role of the VTA-AcbSh circuit in responding for a natural reward was established using non-linear regression (GraphPad Prism 10 Software, Inc.) and GLMM (SAS/STAT Software 9.4). Rodents had to record at least five responses to be included in the analysis necessitating the removal of one male DREADD rat from the 0.1 mg/kg C21 dose. In addition, one male DREADD rat was not tested at the 0.1 mg/kg C21 dose. The number of sucrose reinforcers earned was the dependent variable of interest. Biological sex and surgery served as between-subjects factors, whereas C21 dose and time served as within-subjects factors.

Non-linear regression (GraphPad Prism 10 Software, Inc.) and GLMM (SAS/STAT Software 9.4) were subsequently implemented to evaluate the role of the VTA-AcbSh circuit in responding for concurrently available natural and drug rewards. Rodents had to record at least five total responses to be included in the analysis, yielding sample sizes of: 0 mg/kg: DREADD Male, *n*=4, DREADD Female: *n*=8, Sham Male, *n*=5, Sham Female, *n*=7; 0.01 mg/kg C21: DREADD Male, *n*=9, DREADD Female: *n*=9, Sham Male, *n*=6, Sham Female, *n*=7; 0.03 mg/kg C21: DREADD Male, *n*=10, DREADD Female: *n*=7, Sham Male, *n*=6, Sham Female, *n*=7; 0.1 mg/kg C21: DREADD Male, *n*=10, DREADD Female: *n*=6, Sham Male, *n*=6, Sham Female, *n*=7; 0.3 mg/kg C21: DREADD Male, *n*=9, DREADD Female: *n*=8, Sham Male, *n*=5, Sham Female, *n*=7. The dependent variable of interest (i.e., number of rewards) was log-transformed for analyses of female animals. Biological sex and surgery served as between-subjects factors, whereas C21 dose, time, and reinforcer served as within-subjects factors.

Complementary analyses, including non-linear regression (GraphPad Prism 10 Software, Inc.) and GLMM (SAS/STAT Software 9.4) were conducted to establish the time frame in which C21, relative to saline, altered reinforced responding. Given the pronounced sex differences observed in responding for concurrently available natural and drug rewards, analyses were conducted independently by biological sex, whereby the dependent variable of interest (i.e., Male: Number of Sucrose Rewards; Female: Number of Cocaine Reinforcers) was transformed (i.e., Male: Square Root Transformation; Females: Squared Transformation).

Furthermore, dose (i.e., Male: 0 mg/kg, 0.03 mg/kg C21; Female: 0 mg/kg, 0.1 mg/kg C21) and time served as within-subjects factors.

Sequential administration of C21 and Sal B, a KORD-based DREADDs agonist, was utilized to verify that the alterations in reinforced responding induced by C21 were a result of DREADDs-mediated changes in the VTA-AcbSh circuit; data from this experiment was analyzed using GLMM (SAS/STAT Software 9.4). Given the pronounced sex differences observed in responding for concurrently available natural and drug rewards, analyses were conducted independently by biological sex. For female analyses, the dependent variable of interest was log-transformed. Within-subjects factors included in the analysis were: injection, time, and reinforcer.

For all GLMM analyses, a PROC GLIMMIX statement with a random intercept, Poisson distribution, and unstructured covariance was utilized to statistically evaluate changes in the number of sucrose reinforcers earned. Least squares means with a Tukey correction were conducted post-hoc.

## RESULTS

### Phase 1: All Fischer F344/N male and female rats successfully acquired sucrose and cocaine self-administration

Male and female Fischer F344/N rodents were trained to self-administer sucrose and cocaine under fixed (Sucrose: 5% w/v unless otherwise specified; Cocaine: 0.2 mg/kg/infusion) and/or progressive (Cocaine: 0.75 mg/kg/infusion) ratio schedules of reinforcement.

Independent of biological sex, all rats met the criterion of 60 reinforcers for three consecutive days (**Figure 2A**) under an FR(1) schedule of reinforcement, thereby establishing the acquisition of sucrose self-administration. All female rats successfully acquired sucrose self-administration within 49 test sessions, whereby the rate of acquisition was well-described by a one-phase association (*R*^2^≥0.98). A prominent rightward shift in the number of days required to acquire sucrose self-administration was observed in male animals (Best Fit Function: One-Phase Association (*R*^2^≥0.98)) supporting a decreased rate of task acquisition relative to their female counterparts (Significant Differences in *K*, [*F*(1, 28)=18.01, *p*≤0.001]). Although 16 male rats successfully acquired sucrose self-administration within 60 test sessions, additional measures were necessary to increase responding for four rodents (Water Restriction, *n*=1; Water Restriction and 10% Sucrose Concentration, *n*=3). Therefore, no data was censored, as all rodents successfully acquired sucrose self-administration.

**Figure 2.**
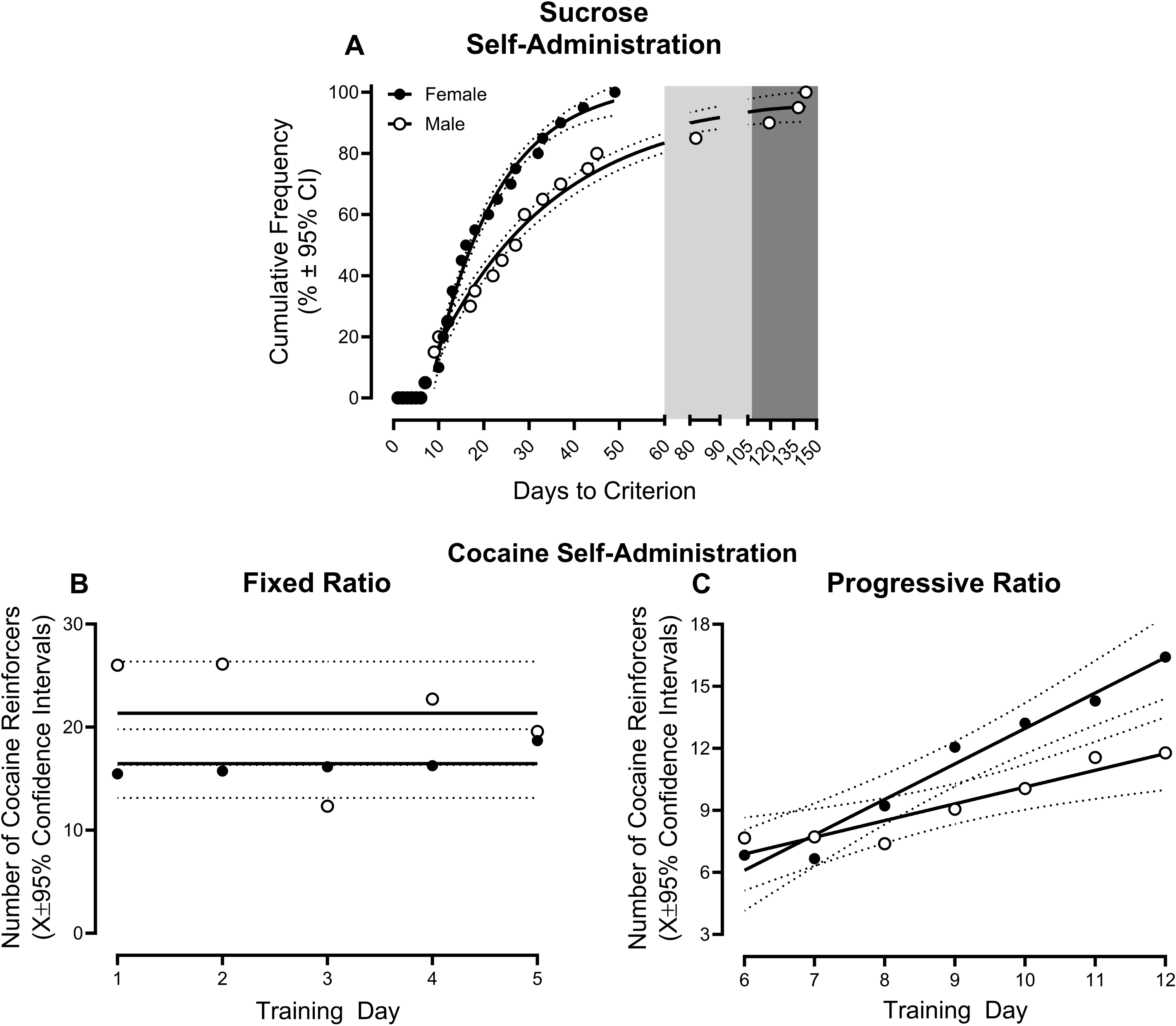
Preliminary Operant Training. Rodents underwent extensive operant self- administration training for sucrose (5% *weight/volume* (*w/v*) unless otherwise specified) or cocaine (Fixed Ratio (FR): 0.2 mg/kg/infusion; Progressive Ratio (PR): 0.75 mg/kg/infusion). (**A**) All animals, independent of biological sex, successfully acquired sucrose self-administration by responding for at least 60 reinforcers for three consecutive days. The light-shaded gray area indicates the time period during which a small proportion of male rodents were water-restricted for up to 18 hours. The dark shaded gray area indicates rats who required an increased sucrose solution concentration (i.e., 10% *w/v* instead of 5% *w/v*) to meet criteria. (**B-C**) Rodents were trained to self-administer cocaine under FR and PR schedules of reinforcement for five and seven days, respectively. Under a PR schedule of reinforcement, male and female animals displayed an escalation of cocaine intake supporting a drug-dependent phenotype. Solid lines represent the best-fit function, whereas dotted lines illustrate the 95% confidence interval (CI).

Rodents acquired cocaine self-administration by responding for a drug reinforcer under both fixed (Days 1-5; **Figure 2B**) and progressive (Days 6-12; **Figure 2C**) ratio schedules of reinforcement. Independent of biological sex, rodents exhibited a prominent linear escalation of cocaine self-administration under a PR schedule of reinforcement (*R*^2^s≥0.88), supporting the development of a drug-dependent phenotype. Nevertheless, the factor of biological sex influenced the rate of cocaine self-administration escalation, whereby female rodents escalated more rapidly than their male counterparts (Biological Sex x Day Interaction, [*F*(1,212)=10.6, *p*≤0.001]; Significant Differences in β_1_, [*F*(1, 246)=6.0, *p*≤0.015]). Results support, therefore, the successful acquisition of both sucrose and cocaine self-administration.

### Phase 2: In male rodents, responding under a single schedule of reinforcement is mediated, at least in part, by neurons in the VTA-AcbSh circuit

After the successful acquisition of sucrose and cocaine self-administration, rodents were intravenously injected with C21 (0, 0.01, 0.03, or 0.1 mg/kg), a DREADDs agonist, to evaluate the role of neurons in the VTA-AcbSh circuit in natural (i.e., sucrose) reward responding.

Indeed, administration of C21, significantly altered responding in a surgery-, sex-, and dose-dependent manner (**Figure 3**; Surgery x Biological Sex x Dose Interaction, [*F*(3, 91)=3.3, *p*≤0.022]). Complementary analyses were conducted independently by sex to establish the locus of the interaction.

**Figure 3.**
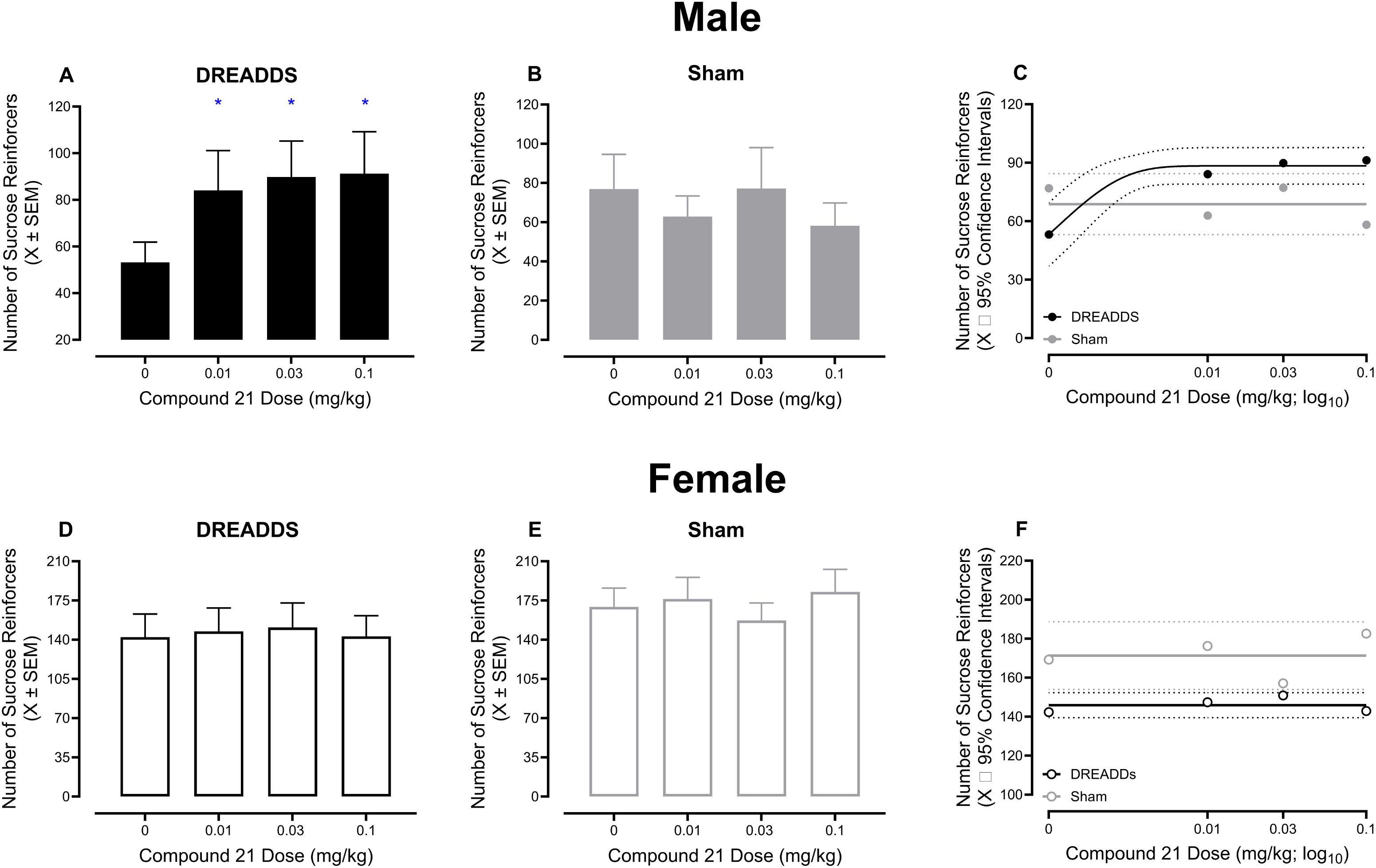
DREADDs Activation Under a Single Schedule of Reinforcement. Animals were treated with varying doses of Compound 21 (C21) in an ascending order to establish how activation of the mesolimbic neurocircuit alters sucrose-reinforced responding under a single schedule of reinforcement. (**A-C**) DREADD, but not sham, male animals exhibited increased sucrose-reinforced responding after administration of 0.01, 0.03, or 0.1 mg/kg C21. (**D-F**) C21- induced activation of the mesolimbic neurocircuit in female animals, independent of DREADD or sham transduction, failed to alter sucrose-reinforced responding under a single schedule of reinforcement. Solid lines represent the best-fit function, whereas dotted lines illustrate the 95% confidence interval (CI). * *p*<0.05

Male rodents transduced with DREADDs exhibited increased responding for a natural reward following pharmacological manipulation of the VTA-AcbSh circuit (**Figure 3A**; Surgery x Dose Interaction, [*F*(3, 46)=4.9, *p*≤0.005]). Specifically, post-hoc comparisons with a Tukey-Kramer correction revealed a statistically significant increase in sucrose-reinforced responding (relative to saline) in male DREADDs, but not sham (*p*>0.05; **Figure 3B**), rodents following 0.01 mg/kg [*t*(46)= -3.2, *p*≤0.05], 0.03 mg/kg [*t*(46)= -3.8, *p*≤0.010], and 0.1 mg/kg [*t*(46)= -3.8, *p*≤0.011] C21 administration. Regression analyses confirmed these findings, whereby a one-phase association (**Figure 3C**; *R*^2^≥0.99) well-described the number of sucrose reinforcers earned by male DREADDs rodents across C21 dose; a horizontal line, however, provided the best-fit function for male sham animals (**Figure 3C**).

Administration of C21 in female rodents transduced with either DREADDs or vehicle, however, failed to alter responding for a natural reward (**Figure 3D-E**; *p*>0.05). Consistently, regression analyses revealed that the number of sucrose reinforcers earned by female DREADDs and sham rodents across C21 dose was well-described by a horizontal line (**Figure 3F**). Collectively, neurons in the VTA-AcbSh circuit appear fundamentally involved in responding for a natural reward in male, but not female, rats.

### Phase 3: Activation of neurons in the VTA-AcbSh circuit alters responding for concurrently available rewards in a reinforcer- and sex-dependent manner

Rodents were subsequently habituated to a concurrent choice self-administration experimental paradigm, whereby responding was reinforced with either 5% sucrose (*w/v*) or 0.2 mg/kg/infusion cocaine. After the establishment of choice behavior, rodents were intravenously injected with C21 (0, 0.01, 0.03, 0.1, or 0.3 mg/kg) to evaluate the role of neurons in the VTA- AcbSh circuit in responding for concurrently available natural (i.e., sucrose) and drug (i.e., cocaine) rewards.

Indeed, administration of C21 immediately prior to the concurrent choice self-administration experimental paradigm significantly altered responding in a surgery-, sex-, reinforcer- and dose-dependent manner (**Figure 4**; Surgery x Biological Sex x Dose x Reinforcer Interaction, [*F*(4, 95)=5.1, *p*≤0.001]). Complementary analyses were conducted independently by sex to establish the locus of the interaction.

**Figure 4.**
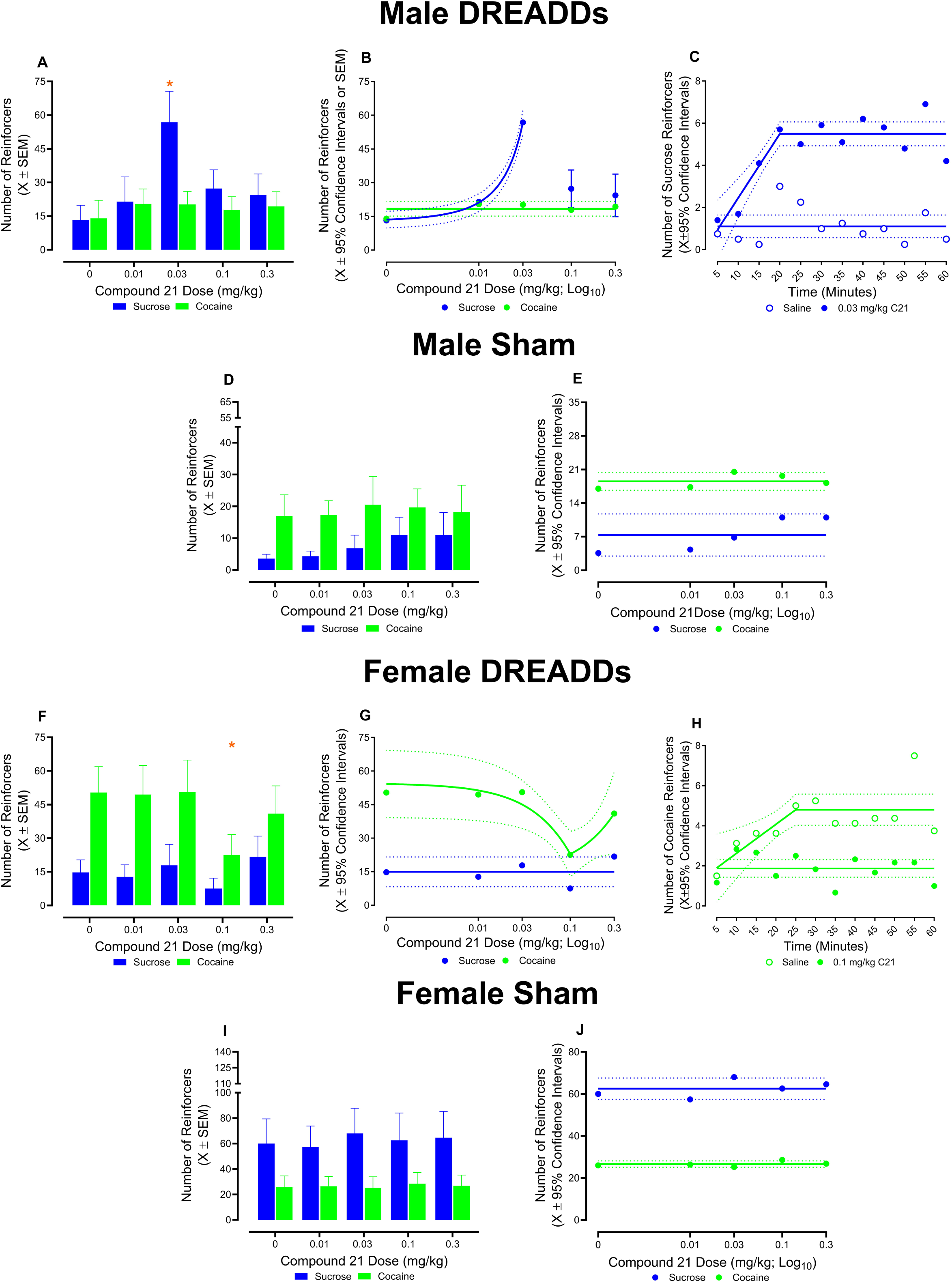
DREADDs Activation Under a Concurrent Choice Self-Administration Experimental Paradigm. Varying doses of Compound 21 (C21) were administered to rodents in an ascending order to evaluate the role of projections from the ventral tegmental area (VTA) to nucleus accumbens (AcbSH) in behavioral allocation during a concurrent choice self- administration experimental paradigm. (**A-C**) Male rodents transduced with DREADDs exhibited increased sucrose-reinforced responding following injection with 0.03 mg/kg C21; C21 administration did not alter responding for cocaine reinforcers at any dose. At the 0.03 mg/kg dose of C21, increased sucrose-reinforced responding began 15 minutes after injection and persisted throughout the 60-minute test session. (**D-E**) C21 treatment did not significantly alter sucrose or cocaine-reinforced responding in sham male animals. (**F-H**) A prominent decrease in cocaine-reinforced responding was observed in female rats transduced with DREADDs after administration of 0.1 mg/kg C21; C21 injection failed to alter sucrose-reinforced responding at any dose. At the 0.1 mg/kg dose of C21, decreased responding for cocaine reinforcers began 15 minutes after treatment and persisted throughout the 60-minute test session. (**I-J**) Neither sucrose nor cocaine-reinforced responding were altered by C21 administration in female sham animals. Solid lines represent the best-fit function, whereas dotted lines illustrate the 95% confidence interval (CI). * *p*<0.05

Pharmacological manipulation of the VTA-AcbSh circuit selectively altered reinforced responding in male rodents in a surgery-, reinforcer-, and dose-dependent manner (**Figure 4A- B**; Surgery x Reinforcer x Dose Interaction, [*F*(4, 45)=7.4, *p*≤0.001]). Specifically, male DREADDs rodents treated with 0.03 mg/kg C21 exhibited a significant increase in sucrose-reinforced responding (Post-Hoc Comparisons with a Tukey-Kramer Correction: DREADDs Saline vs. DREADDs 0.03 mg/kg C21, [*t*(45)= -7.5, *p*≤0.001]). Independent of dose, C21 failed to alter cocaine-reinforced responding in male DREADDs rats (*p*>0.05). Findings were confirmed using linear regression analyses, whereby, in male DREADDs animals, the number of sucrose reinforcers earned from the 0 mg/kg through 0.03 mg/kg C21 dose was well-described by an exponential growth equation (*R*^2^≥0.99); a horizontal line provided the best-fit function for the number of cocaine reinforcers earned. No alterations in natural or drug-reinforced responding were observed in male sham animals following administration of C21 (*p*>0.05; Best-Fit Function for Sucrose or Cocaine Reinforcers: Horizontal Line; **Figure 4D-E**).

Complementary analyses were conducted to elucidate the timeframe in which injection with 0.03 mg/kg C21, relative to saline, increased sucrose-reinforced responding in DREADDs male animals. Indeed, a statistically significant Time x Dose interaction [*F*(11,110)=1.9, *p*≤0.05] supports that the effects of 0.03 mg/kg C21 occurred in a time-dependent manner (**Figure 4C**). Male DREADDs rodents treated with 0.03 mg/kg C21 exhibited a significant increase in sucrose-reinforced responding beginning 15 min after injection; an effect which persisted through 60 min (Post-Hoc Comparisons with a Tukey-Kramer Correction: DREADDs Saline vs. DREADDs 0.03 mg/kg C21, *p*≤0.05; Individual Comparisons Presented in **Supplementary Table 1**). Linear regression analyses verified these observations, whereby a segmental linear regression well-described the number of sucrose reinforcers earned by DREADDs male animals following administration of 0.03 mg/kg C21 (*R*^2^≥0.80). In sharp contrast, the number of sucrose reinforcers earned by DREADDs male animals after treatment with saline remained static throughout the 60-min test session (Best Fit Function: Horizontal Line).

In female animals, IV administration of C21 also selectively altered reinforced responding in a surgery-, reinforcer-, and dose-dependent manner (**Figure 4F-G**; Surgery x Reinforcer x Dose Interaction, [*F*(4, 52)=2.6, *p*≤0.05]). Specifically, female DREADDs rodents treated with 0.1 mg/kg C21 exhibited a significant decrease in reinforced responding for a drug reward (Post-Hoc Comparisons with a Tukey-Kramer Correction: DREADDs Saline vs. DREADDs 0.1 mg/kg C21, [*t*(52)= 6.56, *p*≤0.001]). Sucrose reinforced responding, however, was not altered by pharmacological manipulation of the VTA- AcbSh circuit in female DREADDs animals. Linear regression analyses were implemented to confirm these findings, whereby, in female DREADDs rats, the number of cocaine reinforcers earned across C21 doses was well-described by a segmental linear regression (*R*^2^≥0.969); a horizontal line provided the best-fit function for the number of sucrose reinforcers earned. Administration of C21 in female vehicle rodents, however, induced no alterations to natural or drug-reinforced responding (*p*>0.05; Best-Fit Function for Sucrose or Cocaine Reinforcers: Horizontal Line; **Figure 4I-J**).

Complementary analyses were conducted to elucidate the timeframe in which treatment with 0.1 mg/kg C21, relative to saline, decreased cocaine-reinforced responding in DREADDs female animals. Notably, the effects of 0.1 mg/kg C21 occurred in a time-dependent manner (**Figure 4H**; Time x Dose Interaction, [*F*(11,77)=31.3, *p*≤0.001]). A significant decrease in number of cocaine reinforcers earned by female DREADDs rodents treated with 0.1 mg/kg C21, relative to saline, was observed beginning 15 min after injection; an effect which persisted through 60 min (Post-Hoc Comparisons with a Tukey-Kramer Correction: DREADDs Saline vs. DREADDs 0.03 mg/kg C21, *p*≤0.05; Individual Comparisons Presented in **Supplementary Table 1**). Findings were confirmed using linear regression analyses, whereby, in female DREADDs animals, the number of cocaine reinforcers earned across time after administration of 0.1 mg/kg C21 remained static throughout the 60-min test session (Best Fit Function: Horizontal Line). In sharp contrast, the number of cocaine reinforcers earned by DREADDs female animals after treatment with saline increased steadily from five to 25 min, at which point the number of reinforcers earned within each bin reached asymptote (Best Fit Function: Segmental Linear Regression; *R*^2^≥0.52).

### Phase 3: Alterations in reinforced responding for concurrently available rewards resulted from DREADDs-mediated changes in the VTA-AcbSh circuit

Lastly, sequential administration of C21 and Sal B, a KORD-based DREADDs agonist, was utilized to verify that the alterations in reinforced responding induced by C21 were a result of DREADDs-mediated changes in the VTA-AcbSh circuit.

Male DREADDs rodents were injected with 0.03 mg/kg C21 followed by 0.15 mg/kg SalB. Indeed, sequential administration of C21 and SalB reversed C21-induced effects on sucrose responding (**Figure 5A**; Injection x Reinforcer Interaction, [*F*(2, 8)=16.3, *p*≤0.002]). Specifically, post-hoc comparisons with a Tukey-Kramer correction revealed that sucrose-reinforced responding in male DREADDs rats following 0.03 mg/kg C21 and 0.15 mg/kg SalB was statistically indistinguishable from that which was observed in these animals following saline administration (*p*>0.05).

**Figure 5.**
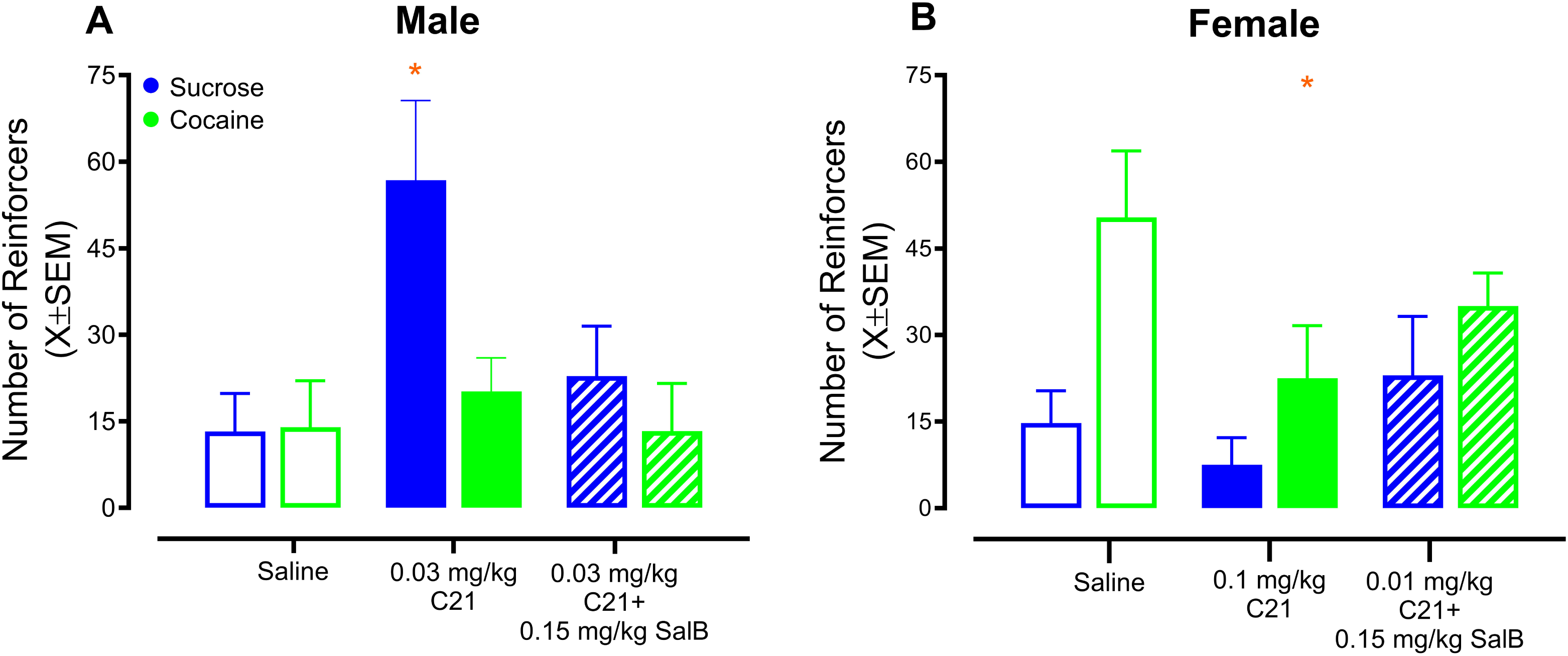
DREADDs Inhibition Under a Concurrent Choice Self-Administration Experimental Paradigm. Rodents were sequentially treated with Compound 21 (C21) and Salvinorin B (SalB) to verify the role of projections from the ventral tegmental area (VTA) to nucleus accumbens (AcbSH) in behavioral allocation. (**A**) DREADD male animals were injected with 0.03 mg/kg C21 followed by 0.15 mg/kg SalB. Although 0.03 mg/kg C21 significantly increased sucrose-reinforced responding, responding following sequential administration of C21 and SalB was statistically indistinguishable from saline. (**B**) DREADD female rodents were injected with 0.1 mg/kg C21 followed by 0.15 mg/kg SalB. Although 0.1 mg/kg C21 significantly decreased responding for a drug (i.e., cocaine) reward, responding following sequential administration of C21 and SalB was statistically indistinguishable from saline. * *p*<0.05 relative to saline

DREADDs female rats were consecutively administered with 0.1 mg/kg C21 and 0.15 mg/kg SalB. Indeed, sequential administration of C21 and SalB reversed C21-induced effects on cocaine-reinforced responding (**Figure 5B**; Injection x Reinforcer Interaction, [*F*(2, 11)=6.5, *p*≤0.012]). Specifically, posthoc comparisons with a Tukey-Kramer correction revealed that cocaine-reinforced responding in female DREADDs rodents following 0.1 mg/kg C21 and 0.15 mg/kg SalB was statistically indistinguishable from that which was observed in these animals following saline administration (*p*>0.05).

### After the completion of behavioral testing, cannula placement and the expression of designer receptors exclusively activated by designer drugs were verified

Cannula placement and DREADDs expression were verified in at least a subset of animals. Green fluorescent protein (GFP) was clearly expressed in the AcbSh indicating appropriate transduction of AAV-CMV-GFP/Cre. Indeed, the majority of the GFP expression was observed in the AcbSh shell (**Supplementary** Figure 1). In rodents transduced with DREADDs, mCherry and mCitrine expression was identified in the parabrachial nucleus of the posterior VTA (**Figure 6B**) corresponding to the infusion of pAAV-hSyn-DIO-hM3D(G_q_)-mCherry and pAAV-hSyn-dF-HA-KORD-IRES-mCitrine, respectively.

## DISCUSSION

Bidirectional control of neurons projecting from the VTA to the AcbSh demonstrates the fundamental role of the mesolimbic neurocircuit in the allocation of behavior. Pharmacological activation of the VTA-AcbSh circuit modulated responding for a natural reward (i.e., sucrose) under a single schedule of reinforcement in a sex-dependent manner. During a concurrent choice self-administration experimental paradigm, C21 administration induced a selective shift in the allocation of behavior in rodents transduced with DREADDs. Specifically, male and female DREADDs animals exhibited a robust increase in responding for a natural reward and a prominent decrease in drug-reinforced (i.e., cocaine) responding, respectively, following pharmacological activation of the VTA-AcbSh circuit. The sequential activation of hM3D(G_q_) and KORD DREADDs within the same neuronal population validated the role of the VTA-AcbSh circuit in reinforced responding for concurrently available natural and drug-reinforced responding. Taken together, the VTA-AcbSh circuit plays an integral role in drug-biased choice affording a key target for the development of novel pharmacotherapies for cocaine use disorder.

A concurrent choice self-administration experimental paradigm was implemented to evaluate the role of the mesolimbic dopamine system in behavioral allocation. In preclinical choice procedures, animals are trained to respond on two different manipulanda (e.g., lever) for two distinct reinforcers (e.g., IV bolus of cocaine or orally available sucrose), thereby requiring an individual to allocate behavior (or “choose”) between concurrently available rewards. The translational relevance of choice procedures cannot be understated, as these procedures model a key aspect of the clinical phenotype of cocaine use disorder: the use of substances in a complex environment with other concurrently available reinforcers. Although choice procedures are being increasingly utilized to evaluate novel therapeutics for the treatment of cocaine use disorder (for review, [42]), they remain underutilized as a tool to investigate the neurobiological mechanisms that contribute to the allocation of behavior.

The validity of DREADDs experiments is dependent, at least in part, on the assumption that their chemical actuator is pharmacologically inert and exhibits high brain bioavailability.

GPCR-based DREADDs (e.g., hM3D(G_q_)) were originally designed to respond exclusively to the synthetic ligand CNO [8], a metabolite of clozapine; CNO, at the time, was considered to be a pharmacologically inert molecule [8,43]. Subsequent studies, however, revealed that CNO binds to endogenous receptors [41,44] and questioned the ability of peripherally administered CNO to penetrate the blood-brain barrier [44]. Indeed, research supports the reverse metabolism of peripherally administered CNO to clozapine, which subsequently crosses the blood-brain barrier and activates DREADDs [44–45]. As a result, a second generation of synthetic DREADD ligands, including C21 [18], were engineered. Although C21 also binds to endogenous ligands [18,41], the novel compound exhibits improved selectivity compared to clozapine [18]. With regards to brain bioavailability, pharmacokinetic experiments demonstrated a graduate increase in mean brain concentrations of C21 from 15 to 60 minutes post-injection [41].

Due to the novelty of C21 as a potent chemical actuator, fewer studies have characterized this DREADD synthetic agonist; an opportunity capitalized upon in the present study via the utilization of both dose- and time-response experimental paradigms.

Unsurprisingly, the route of administration influences the most efficacious dose of C21, whereby, in prior studies, robust electrophysiological [46] and behavioral [47] responses have been reported in DREADDs rodents intraperitoneally injected with 0.5 to 10 mg/kg C21. Herein, prominent behavioral effects were induced by significantly lower doses of intravenously administered C21 (i.e., 0.01 to 0.1 mg/kg), as IV administration bypasses the absorption process completely. Utilization of a dose-response experimental paradigm also revealed profound sex differences in optimized C21 dosages for males and females, an effect which may result from sex differences in GPCR pharmacodynamics (for review, [48]) or ligand pharmacokinetics (for review, [49]). Furthermore, the data were deconstructed to evaluate time-dependent behavioral responses, whereby IV C21, independent of biological sex, induced profound behavioral effects from 15 to 60 minutes post-injection; the onset of action of IV C21 is consistent with pharmacokinetic studies [41] and behavioral responses following intraperitoneal injections of C21 [19]. A thorough characterization of DREADD ligands, including C21, is integral to establishing the advantages and constraints of rapidly evolving chemogenetics techniques.

KORD-based DREADDs, in contrast, utilize a unique chemical actuator (i.e., SalB) affording a fundamental opportunity to bidirectionally control neuronal activity in the same animal. SalB, a metabolite of the KOR selective agonist salvinorin A (SalA), is a pharmacologically inert ligand, as it exhibits minimal, if any, affinity for KOR, other endogenous receptors, or GPCR-based DREADDs [21–22, 50]. From a pharmacokinetics perspective, SalB, when administered intravenously, rapidly enters and clears the brain [23]. Both C21 and SalB, therefore, represent significantly improved, albeit imperfect, DREADDs synthetic agonists for GPCR- and KORD-based DREADDs.

Given potential limitations in the currently available DREADD chemical actuators, experimental design considerations were integral to conducting an interpretable behavioral study. First, the current study implemented stringent controls. Four appropriately powered experimental groups, including male DREADDs, male sham (i.e., DREADD-free), female DREADDs, and female sham were intravenously injected with the same doses of C21; the inclusion of a non-DREADD-expressing control group is in accordance with recommendations for well-controlled DREADD experiments [51–53]. Indeed, C21 failed to alter reinforced- responding in sham animals, thereby demonstrating that behavioral alterations result from the pharmacological manipulation of the VTA-AcbSh circuit. Second, results were verified via the bidirectional control (i.e., C21 + SalB) of neuronal activity in the mesolimbic neurocircuit. When 0.15 mg/kg SalB was intravenously administered 15 minutes after C21, the prominent C21- induced increase in sucrose-reinforced responding and decrease in cocaine-reinforced responding observed in male and female DREADD animals, respectively, was mitigated. Utilization of a concurrent choice self-administration experimental paradigm, in combination with a rigorous experimental design, affords a fundamental opportunity to infer the role of the mesolimbic neurocircuit in the allocation of behavior.

Establishing the role of the VTA-AcbSh circuit in drug-biased choice enhances our understanding of the neural circuits in cocaine-related behaviors. Indeed, prior work has revealed several nuanced, pathway-specific roles for dopaminergic VTA neurons in cocaine-seeking behaviors [54] and cocaine potency [55]. Specifically, G_q_-stimulation of VTA DA neurons induced cocaine relapse [54] and increased cocaine potency [55]; G_i_ signaling, in sharp contrast, selectively reduced cocaine reinstatement behavior [54] and decreased cocaine potency [55]. Chemogenetic approaches have also been instrumental in deconstructing the role of other brain regions (e.g., Lateral Orbitofrontal Cortex; [56]) and neural circuits (e.g., Ventral Pallidum-VTA; [57]; Ventromedial Prefrontal Cortex- AcbSh Shell, [58]) in key behavioral features of cocaine use, including risk-taking behavior [56], cocaine-seeking [57] and cue-induced reinstatement [58]. Taken together, dopaminergic VTA neurons may serve as a key therapeutic target for the development of novel pharmacotherapies, as they are involved not only in drug-biased choice, but also in other cocaine-related behaviors.

Under homeostatic conditions (i.e., 0 mg/kg C21), male animals allocated behavior equally to natural and drug rewards; female rodents, in sharp contrast, exhibited a clear preference for drug rewards. Activation of the VTA-AcbSh circuit mitigated drug-biased choice via increased responding for natural rewards and decreased drug-reinforced responding in male and female rodents, respectively. Albeit, neither male nor female rodents achieved total abstinence, an undeniably high bar [59], following activation of the mesolimbic neurocircuit; the abstinence endpoint, however, has precluded the development of a U.S. Food and Drug Administration-approved medication for cocaine use disorder. Considering these challenges, experts (e.g., [60–61]) and individuals with lived experience [62] are advocating for the development of clinically meaningful non-abstinent outcomes, including decreased stimulant use, as observed in the present study. Indeed, in clinical populations, reduced stimulant use was significantly associated with decreased drug-seeking behaviors and improved psychological and psychosocial functioning [63–64].

Collectively, the fundamental role of the mesolimbic dopamine system in behavioral allocation was demonstrated via the activation and inhibition of DREADDs in a translationally relevant concurrent choice self-administration experimental paradigm. Indeed, activation of the VTA-AcbSh circuit mitigates drug-biased choice in a clinically meaningful manner. Establishing the neural circuits that specify behavioral allocation using a rigorous experimental design and translationally relevant behavioral tasks may provide a key target for the development of novel pharmacotherapies.

## AUTHOR CONTRIBUTIONS

**Kristen A. McLaurin:** Writing-original draft, Writing-review & editing, Formal analysis. **Jessica M. Illenberger:** Writing-review & editing, Data curation. **Hailong Li:** Writing-review & editing, Data curation. **Rosemarie M. Booze:** Writing-review & editing, Funding acquisition, Conceptualization. **Charles F. Mactutus:** Writing-review & editing, Funding acquisition, Conceptualization.

