## Supplementary Information for "SEX-DEPENDENT MODULATION OF BEHAVIORAL ALLOCATION VIA VENTRAL TEGMENTAL AREA-NUCLEUS ACCUMBENS SHELL CIRCUITRY"

789 South Limestone Street

Lexington, KY 40508

^b^Cognitive and Neural Science Program

Department of Psychology

Barnwell College

University of South Carolina

1512 Pendleton Street

Columbia, SC 29208

* These authors contributed equally.

**Word Count:** 6423

**Number of Tables, Figures:** 0, 6

**Abbreviated Title (≤50 characters):** Modulation of Concurrent Choice

**Address Proofs and Correspondence To:**

Charles F. Mactutus, Ph.D.

Professor and Fellow AAAS

Department of Psychology

1512 Pendleton Street

University of South Carolina

Columbia, SC 29208

**Table 1: Post-Hoc Comparisons of the Statistically Significant Dose x Time Interaction.** After establishing the statistically significant Sex x Dose x Time interaction in the overall analyses, complementary analyses, including post-hoc comparisons, were conducted independently by biological sex. *The *p* value reported in the Statistical Output columns utilized a Bonferroni correction for multiple comparisons.

|  | | | | **Male** | **Female** |
| --- | --- | --- | --- | --- | --- |
| **Dose** | **Time** | **Dose** | **Time** | **Statistical Output*** | |
| Saline (0 mg/kg) | 7 | 0.01 mg/kg C21 | 7 | *t*(140)= -3.16, *p*=0.540 | *t*(135)= 0.56, *p*=1.000 |
| Saline (0 mg/kg) | 7 | 0.03 mg/kg C21 | 7 | *t*(140)= -3.79, *p*=0.061 | *t*(135)= 3.37, *p*=0.268 |
| Saline (0 mg/kg) | 7 | 0.1 mg/kg C21 | 7 | *t*(140)= -3.14, *p*=0.566 | *t*(135)= 2.32, *p*=1.000 |
| Saline (0 mg/kg) | 14 | 0.01 mg/kg C21 | 14 | *t*(140)= -1.03, *p*=1.000 | *t*(135)= 1.39, *p*=1.000 |
| Saline (0 mg/kg) | 14 | 0.03 mg/kg C21 | 14 | *t*(140)= -0.33, *p*=1.000 | *t*(135)= -1.75, *p*=1.000 |
| Saline (0 mg/kg) | 14 | 0.1 mg/kg C21 | 14 | *t*(140)= -1.63, *p*=1.000 | *t*(135)= -1.51, *p*=1.000 |
| Saline (0 mg/kg) | 21 | 0.01 mg/kg C21 | 21 | ***t*(140)= -4.86, *p*=0.001** | *t*(135)= -0.64, *p*=1.000 |
| Saline (0 mg/kg) | 21 | 0.03 mg/kg C21 | 21 | *t*(140)= -3.59, *p*=0.124 | *t*(135)= -1.26, *p*=1.000 |
| Saline (0 mg/kg) | 21 | 0.1 mg/kg C21 | 21 | ***t*(140)= -6.03, *p*=0.001** | *t*(135)= -2.11, *p*=1.000 |
| Saline (0 mg/kg) | 28 | 0.01 mg/kg C21 | 28 | *t*(140)= -3.82, *p*=0.055 | ***t*(135)= -3.98, *p*=0.031** |
| Saline (0 mg/kg) | 28 | 0.03 mg/kg C21 | 28 | ***t*(140)= -8.93, *p*=0.001** | ***t*(135)= -4.15, *p*=0.016** |
| Saline (0 mg/kg) | 28 | 0.1 mg/kg C21 | 28 | ***t*(140)= -6.85, *p*=0.001** | *t*(135)= -2.03, *p*=1.000 |
| Saline (0 mg/kg) | 35 | 0.01 mg/kg C21 | 35 | ***t*(140)= -6.39, *p*=0.001** | *t*(135)= 1.40, *p*=1.000 |
| Saline (0 mg/kg) | 35 | 0.03 mg/kg C21 | 35 | ***t*(140)= -6.68, *p*=0.001** | *t*(135)= -1.45, *p*=1.000 |
| Saline (0 mg/kg) | 35 | 0.1 mg/kg C21 | 35 | ***t*(140)= -7.51, *p*=0.001** | *t*(135)= 2.31, *p*=1.000 |
| Saline (0 mg/kg) | 42 | 0.01 mg/kg C21 | 42 | *t*(140)= -3.46, *p*=0.201 | *t*(135)= -1.74, *p*=1.000 |
| Saline (0 mg/kg) | 42 | 0.03 mg/kg C21 | 42 | *t*(140)= -1.31, *p*=1.000 | *t*(135)= 0.37, *p*=1.000 |
| Saline (0 mg/kg) | 42 | 0.1 mg/kg C21 | 42 | *t*(140)= -0.30, *p*=1.000 | *t*(135)= 0.69, *p*=1.000 |

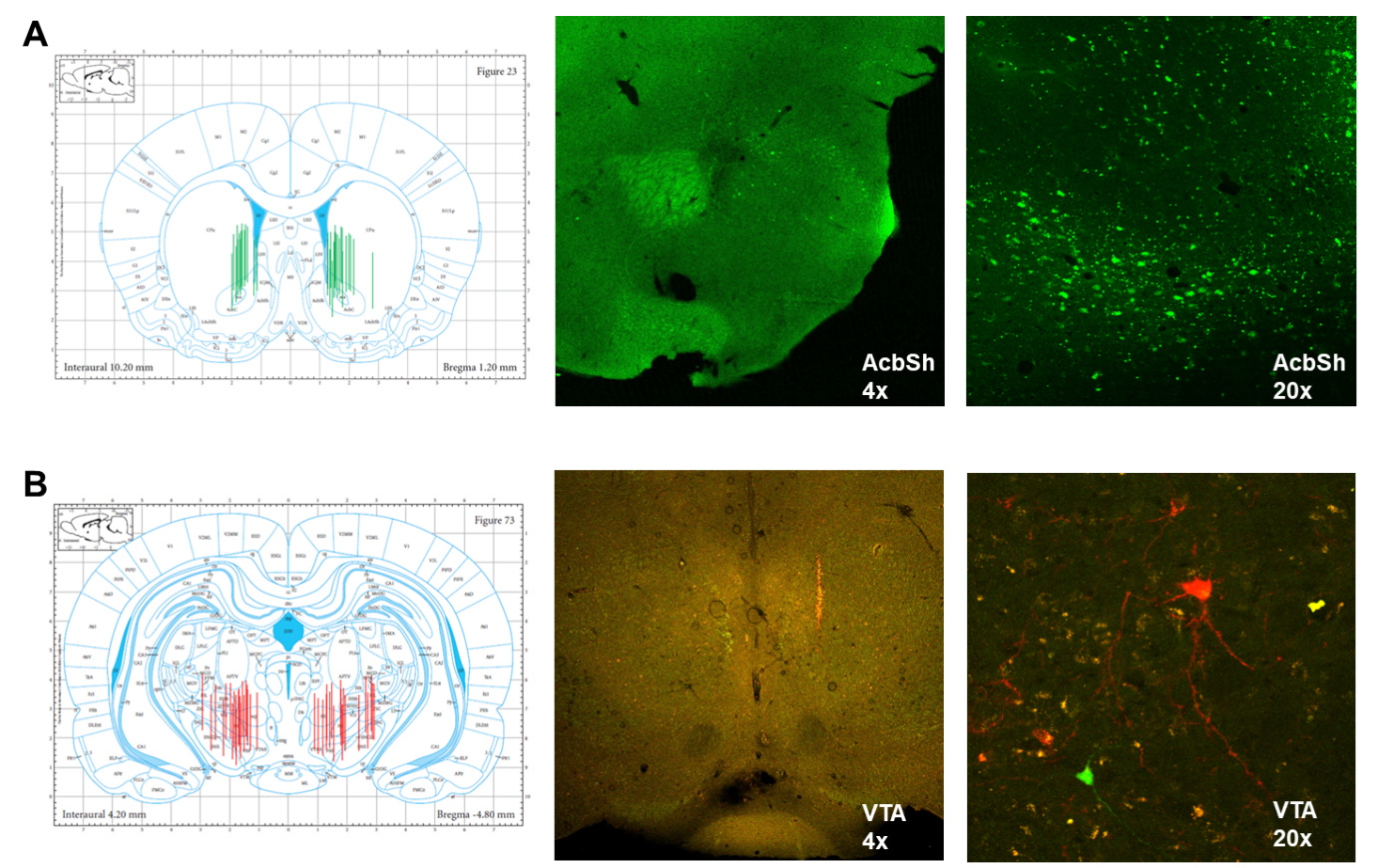

**Supplementary Figure 1. Verification of Cannula Placement and DREADDs Expression.** (**A**) Green fluorescent protein (GFP) was clearly expressed in the nucleus accumbens (AcbSh), whereby the majority of the expression was observed in the AcbSh shell. (**B**) In DREADDs rodents, mCherry and mCitrine expression was identified in the parabrachial nucleus of the posterior ventral tegmental area (VTA).
